## Supplemental Figures for "Learning from Synthetic Dataset for Crop Seed Instance Segmentation"

### Supplemental Materials

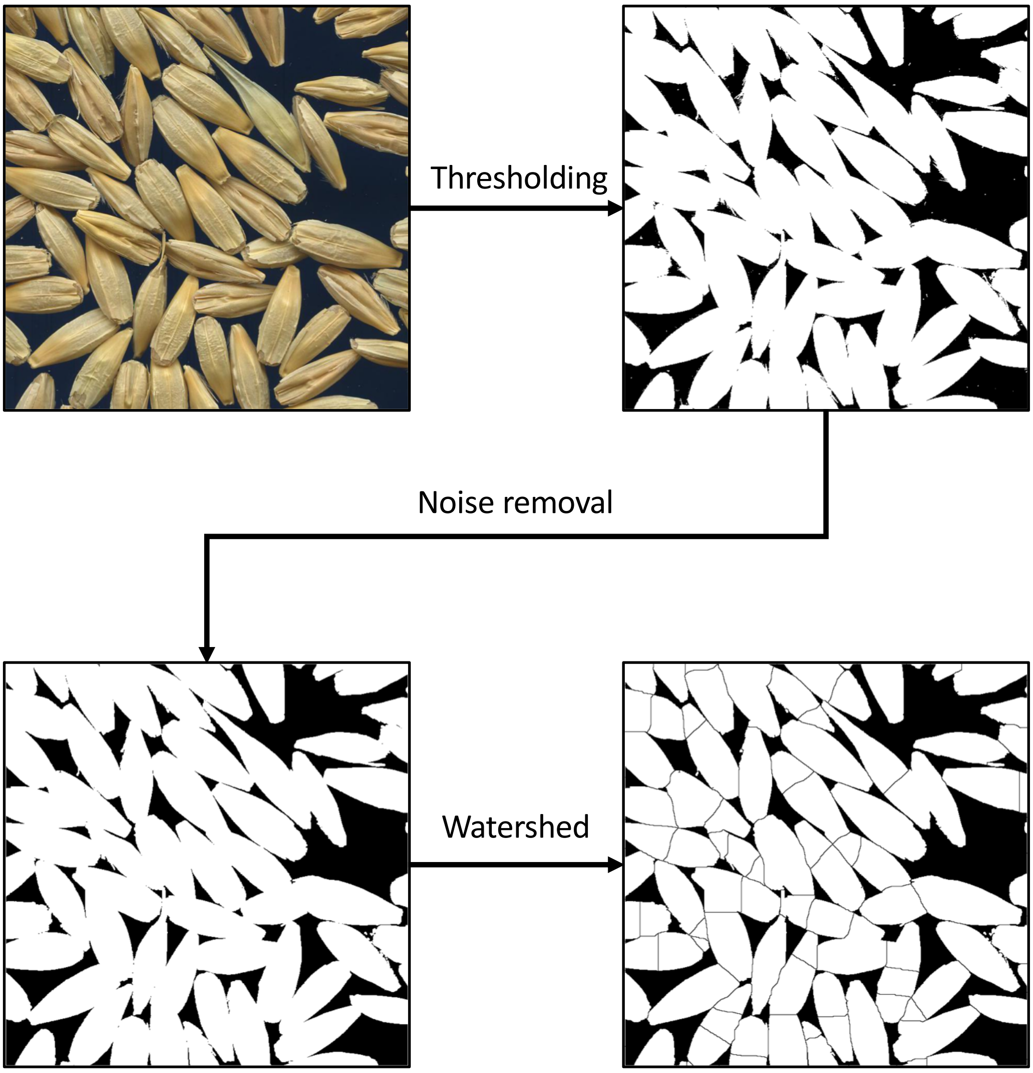


**Fig. S1 | Example of an attempt to segment seed regions by a conventional image analysis method.** Thresholding and noise removal can isolate the seed area from background, but previously known segmentation methods (e.g., watershed) fails to isolate single seed regions.

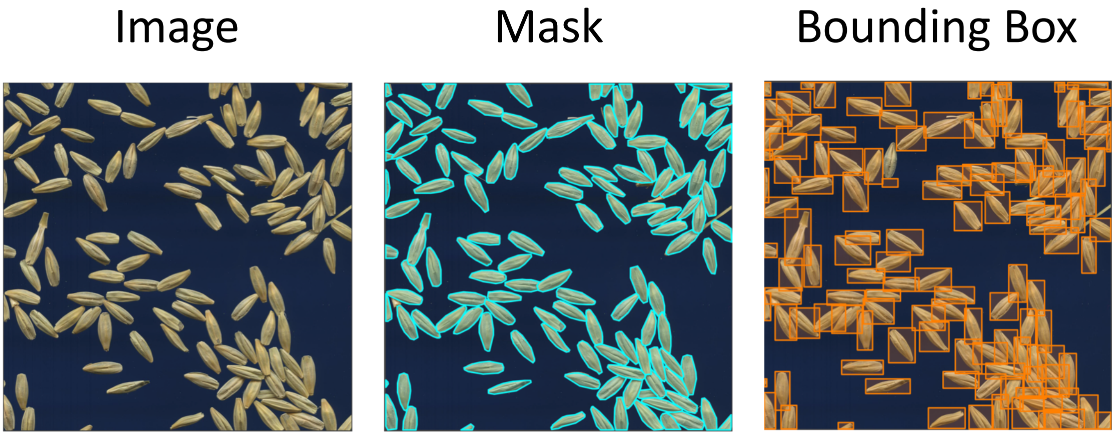


**Fig. S2 | Example of a manual annotation for creating a training data.** A bounding box coordinate and object mask regions are required for training an instance segmentation neural network, which are labor intensive process.

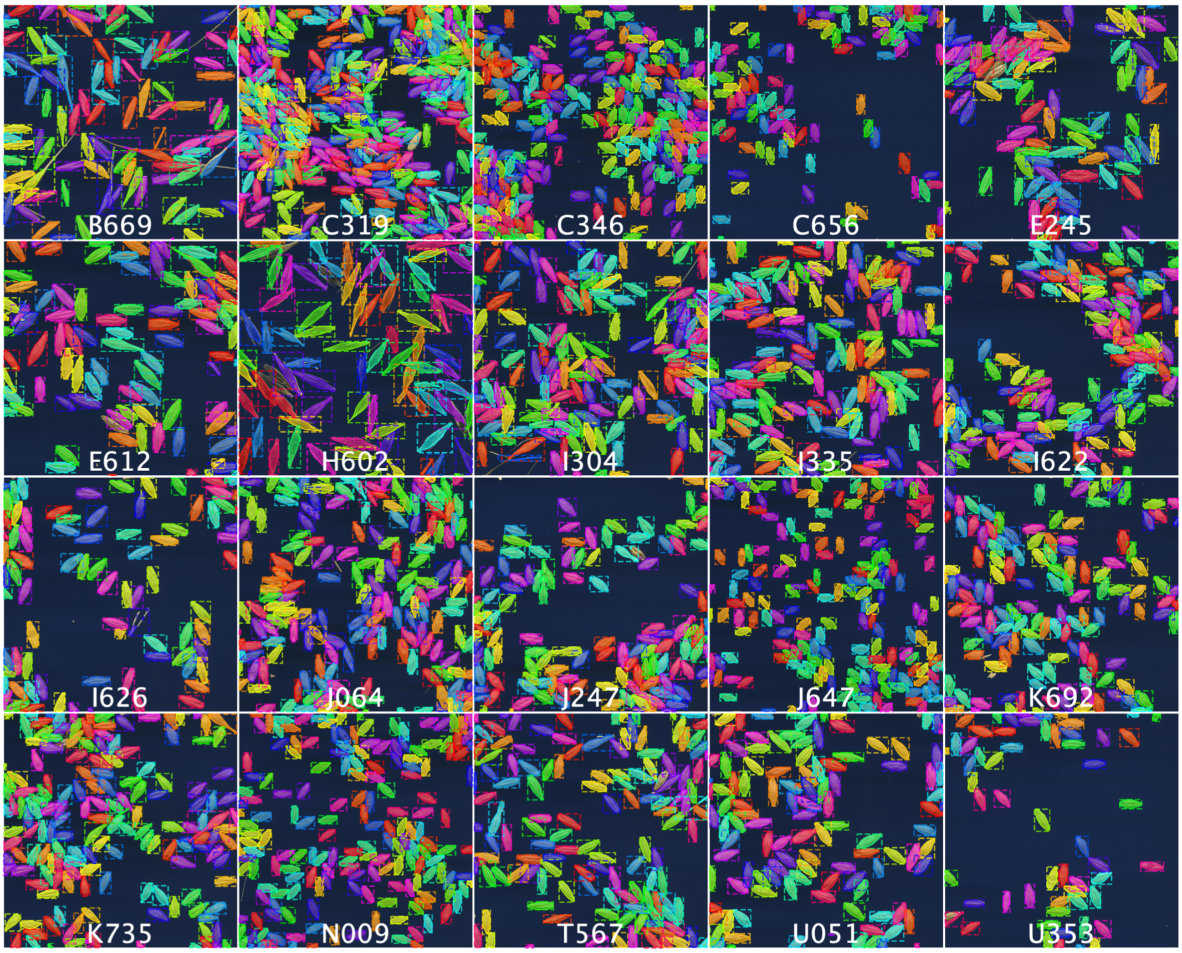


**Fig. S3 | All graphical output of the real-world test data annotated by a trained Mask-RCNN.**

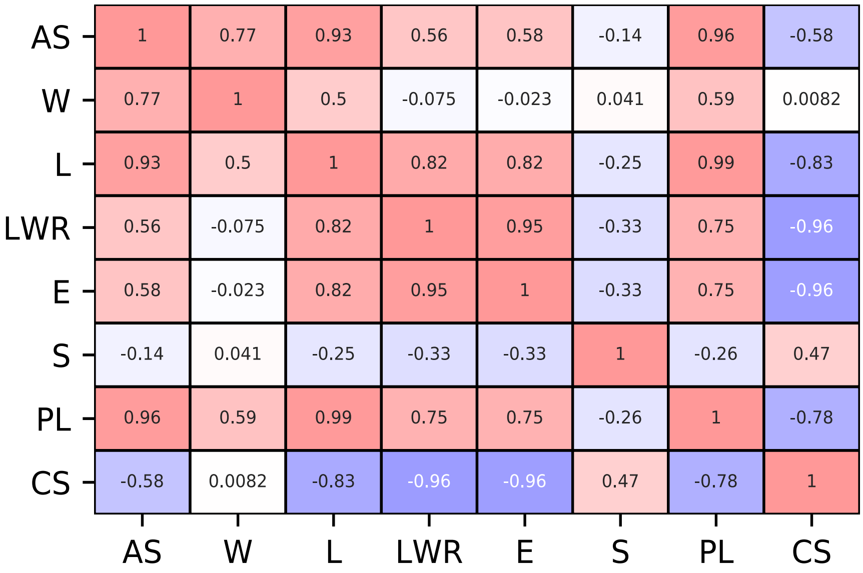


**Fig. S4 | Pearson correlation coefficient matrix of 8 morphological descriptors.**

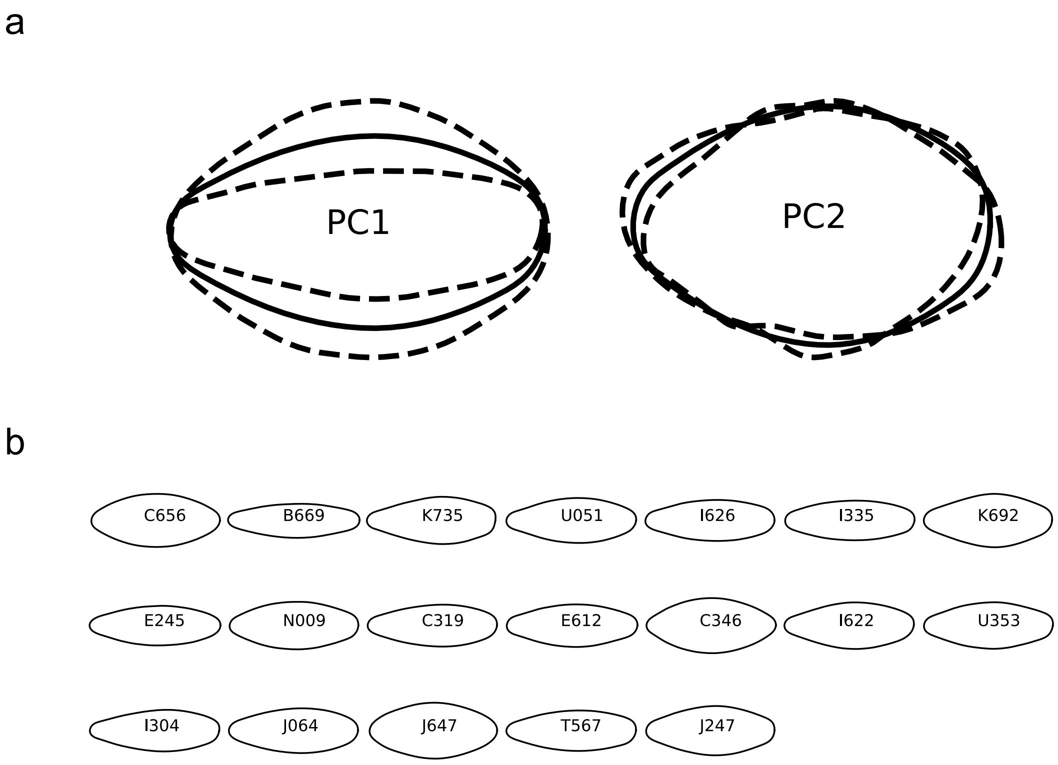


**Fig. S5 | Reconstruction of contours by Elliptic Fourier descriptors (EFDs).** (a) Variation of seed shape that can be accounted for the principal component 1 (PC1) and principal component 2 (PC2). Contours were reconstructed from the corresponding principal component and equal to mean (solid line) or two times the standard deviation (dashed lines). (b) A representative contour of seed shape of respective cultivars reconstructed by the mean values of EFDs coefficients.
